# BMP signalling directs a fibroblast-to-myoblast conversion at the connective tissue/muscle interface to pattern limb muscles

**DOI:** 10.1101/2020.07.20.211342

**Authors:** Joana Esteves de Lima, Cédrine Blavet, Marie-Ange Bonnin, Estelle Hirsinger, Glenda Comai, Laurent Yvernogeau, Léa Bellenger, Sébastien Mella, Sonya Nassari, Catherine Robin, Ronen Schweitzer, Claire Fournier-Thibault, Shahragim Tajbakhsh, Frédéric Relaix, Delphine Duprez

## Abstract

Positional information driving limb muscle patterning is contained in lateral plate mesoderm-derived tissues, such as tendon or muscle connective tissue but not in myogenic cells themselves. The long-standing consensus is that myogenic cells originate from the somitic mesoderm, while connective tissue fibroblasts originate from the lateral plate mesoderm. We challenged this model using cell and genetic lineage tracing experiments in birds and mice, respectively, and identified a subpopulation of myogenic cells at the muscle tips close to tendons originating from the lateral plate mesoderm and derived from connective tissue gene lineages. Analysis of single-cell RNA-sequencing data obtained from limb cells at successive developmental stages revealed a subpopulation of cells displaying a dual muscle and connective tissue signature, in addition to independent muscle and connective tissue populations. Active BMP signalling was detected in this junctional cell sub-population and at the tendon/muscle interface in developing limbs. BMP gain- and loss-of-function experiments performed *in vivo* and *in vitro* showed that this signalling pathway regulated a fibroblast-to-myoblast conversion. We propose that localised BMP signalling converts a subset of lateral plate mesoderm-derived fibroblasts to a myogenic fate and establishes a boundary of fibroblast-derived myonuclei at the muscle/tendon interface to control the muscle pattern during limb development.

## Introduction

Skeletal muscle patterning is a developmental process that controls the formation of each muscle at the right position and time, leading to the highly complex organisation of over 600 individual skeletal muscles in human, each with a specific shape, size, innervation and attachment to the skeletal system. The cellular and molecular mechanisms that regulate limb muscle patterning remain poorly understood. The current consensus emerging from experimental embryology in birds is that limb myogenic cells are of somitic mesoderm, while connective tissue (CT) fibroblasts of muscle attachments originate from the lateral plate mesoderm^1–4^. Consistently, genetic lineage tracing experiments in mice have shown that developmental PAX7+ muscle progenitors originate from the *Pax3* lineage in limbs^5–7^ and that muscle attachments originate from CT fibroblast lineages^8,9^. Previous embryological surgical experiments in chicken embryos demonstrated that the positional information for limb muscles and associated innervation (motoneuron axons) is contained in the lateral plate mesoderm and not in somitic-derived tissues^4,10–14^. This positional information for muscle patterning is maintained over time in lateral plate mesoderm-derived tissues, such as muscle CT^15^ and tendons^16^. The molecular signals produced from lateral plate-derived tissues that drive the correct positioning of limb muscles during development are not fully identified. To date, the TBX4/5, TCF4 and OSR1 transcription factors expressed in limb CT fibroblasts are recognized to act in a non-cell autonomous manner to regulate limb muscle patterning^9,17,18^. While it is possible that CT fibroblasts and myogenic cells interact through secreted signals at the CT/muscle interface, the precise nature and mechanisms of these interactions are currently unknown. The CT/muscle interface is likely to be the place where muscle patterning occurs and one would expect that these junctional cells show specificities in relation to their interface status. The myotendinous junction is a CT/muscle interface that links muscle to tendon and is fully formed at postnatal stages^19^. The mechanisms driving the establishment of the myotendinous junction during development are poorly studied. There is a regionalisation of patterning signals at the tendon/muscle interface, since known signalling molecules display a regionalised expression in a subset of myonuclei at muscle tips close to tendons^20–23^; however, the functions of these regionalized genes in the establishment of the myotendinous junction are not understood.

Here we identify a fibroblast-to-myoblast conversion that takes place at the CT/muscle interface under the control of regionalised BMP signalling to pattern limb muscles.

## Results

To identify the cell origin operating at the muscle/CT interface in developing limbs, we first revisited the embryological origins of CT and myogenic cell populations using cell lineage experiments in avian embryos and genetic lineage tracing experiments in mice. As previously demonstrated^1,2^, isotopic/isochronic quail-into-chicken presomitic mesoderm grafts performed at the presumptive limb regions showed that the vast majority of muscle cells derive from the somitic mesoderm (Fig. 1a-d, Extended Data Fig. 1). However, careful examination of these grafts also identified a subpopulation of myonuclei (nuclei of multinucleated myotubes) that did not have a somitic origin. These myonuclei were located at the extremities of myotubes, close to tendons visualized with collagen XII (Fig. 1a-e, Extended Data Fig. 1). Similarly, a subset of MYOG+ myoblasts (Fig. 1f,g) and PAX7+ muscle progenitors (Extended Data Fig. 2) located at muscle tips were also not of somitic origin. We conclude that, in contrast to our current understanding, a small fraction of PAX7+ muscle progenitors, MYOG+ myoblasts and myonuclei are not derived from somites. To determine from which embryological regions these non-somitic-derived muscle cells originate, we performed the converse experiments and replaced the limb lateral plate of chicken embryos by the quail equivalent region. Consistent with classical lineage descriptions^1,2,4^, this type of graft showed the lateral plate mesoderm origin of cartilage, tendon and muscle connective tissue (Extended Data Fig. 3). However, we also found myonuclei of lateral plate mesoderm origin at muscle tips in close vicinity to tendons visualized with *SCX* expression (Fig. 1h-k). These grafting experiments in avian embryos show an unexpected contribution of lateral plate mesoderm-derived cells to the myogenic lineage at the muscle tips close to tendons in limbs.

**Figure 1.**
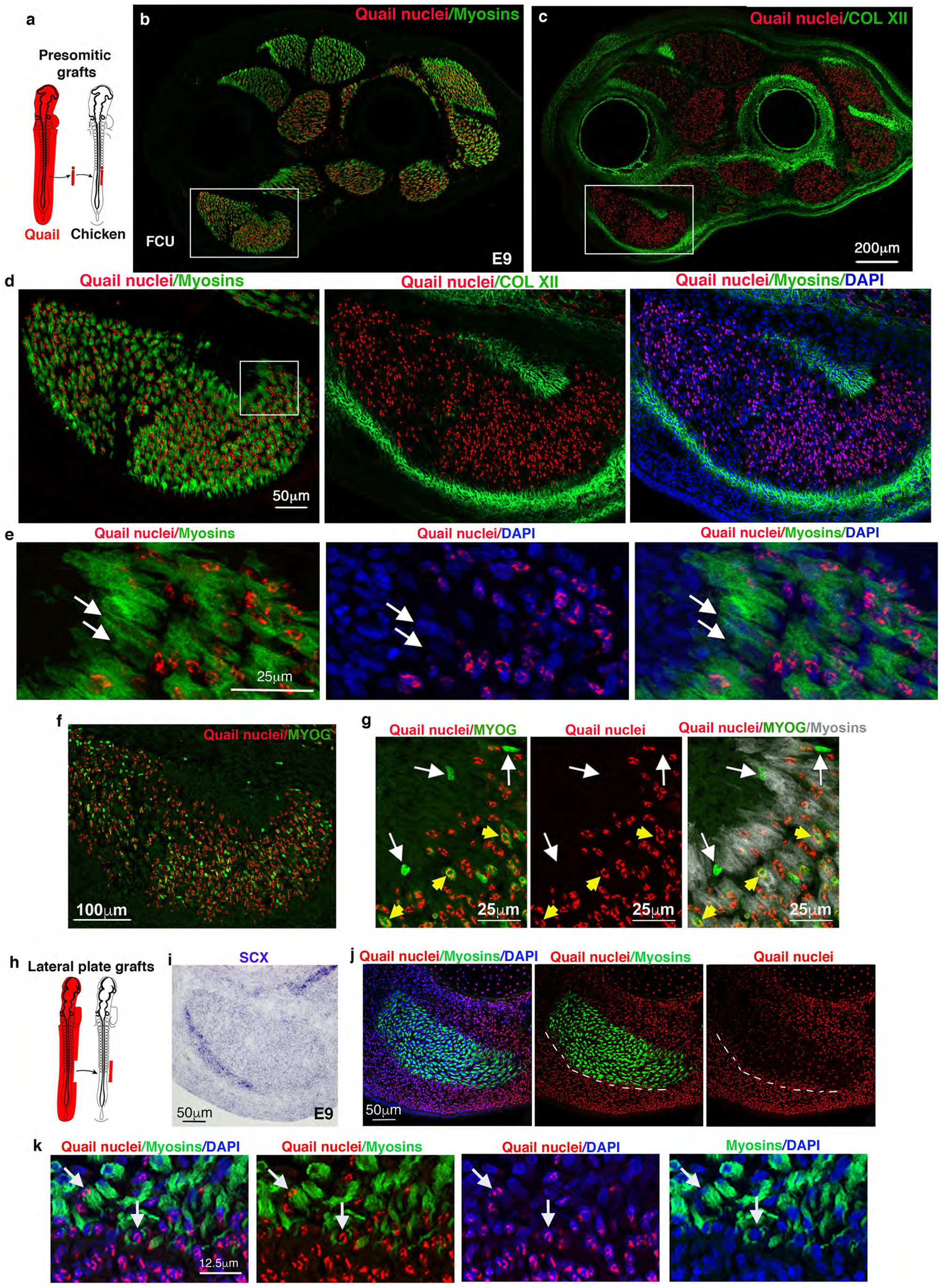
A subpopulation of muscle cells do not originate from somites but from lateral plate in chicken limbs. (**a-g**) Limb muscle organisation after quail-into-chicken presomitic mesoderm grafts. (**a**) Schematic of limb presomitic mesoderm grafts. (**b,c**) Transverse and adjacent limb sections of E9 presomitic mesoderm grafted-embryos were immunostained with QCPN (quail nuclei) and MF20 (myosins) (b), and with QCPN (quail nuclei) and ColXII (tendons) antibodies (c). (**d**) High magnification of FCU, a ventral and posterior muscle immunostained with QCPN (quail nuclei), MF20 (myosins) and ColXII (tendons) antibodies combined with DAPI (nuclei). (**e**) High magnification of muscle tips (close to tendons), squared in (d), showing non-quail nuclei within myosin+ cells (arrows). (**f**) High magnification of FCU immunostained with QCPN (quail nuclei) and MYOG antibodies. (**g**) Focus on muscle tip regions showing MYOG+ nuclei that are not quail+ (white arrows). Yellow arrowheads point to MYOG+ that are quail+. (**h-k**) Limb muscle organisation after quail-into-chicken lateral-plate mesoderm grafts. (h) Schematic of limb lateral plate mesoderm grafts. (**i,j**) Transverse and adjacent limb sections were hybridised with SCX probe (blue) to label tendons (i) and immunostained with QCPN (quail antibody), MF20 (myosins) combined with DAPI, focused on the FCU (j). (**k**) High magnification of muscle tip regions close to tendons showing the high density of quail nuclei of lateral plate mesoderm origin. Arrows point to quail myonuclei in myosin+ cells.

To assess if the unexpected lateral plate mesoderm contribution to limb muscle in chicken is conserved in mice, we analysed PAX7 expression within the *Pax3* lineage. In contrast to the model supporting that *Pax3*+somitic cells give rise to all PAX7+ cells in mice^5–7^, we observed sparse PAX7+ cells that were not *Pax3*-derived (Tomato-negative cells), with a preferential location close to tendons at foetal stages (Fig. 2a-c, Extended Data Fig. 4a). To determine the developmental origin of this subpopulation of non-*Pαx3*-derived PAX7+cells, we performed genetic lineage tracing analyses with the limb CT fibroblast markers *Scx* and *Osr1*. The bHLH transcription factor, SCX is a recognized tendon cell marker during development, which is involved in tendon formation^24,25^. Using the *Scx^Cre^:R26^stop/Tom^* mice^8,26^, we identified PAX7+ and MYOD/MYOG+ myogenic cells within the *Scx* lineage in limbs of E12.5 and E14.5 embryos (Fig. 2d,e, Extended Data Fig. 4b-e). These myogenic cells of CT fibroblast origin were preferentially found surrounding muscle masses at E12.5 and were concentrated close to tendons at E14.5 in mouse limbs (Fig. 2d,e, Extended Data Fig. 4b-e). The zinc finger transcription factor OSR1 is a key regulator of irregular CT differentiation^9^. We performed lineage tracing using *Osr1^CreERT2^* as a driver^9,27,28^ combined with the *Pax7* reporter mouse (*Pax7^GFP-Puro-nlacZ^* = *Pax7^GPL^*), a reporter gene that is not expressed in CT cells, in order to provide a more stringent lineage analysis and circumvent some specificity problems associated with lineage tracing experiments. Using this combination, we identified a subpopulation of PAX7+ muscle progenitors, MYOD/MYOG+ myoblasts and myonuclei that were nlacZ+ (*Osr1* origin) in E15.5 mouse limbs (Fig. 2f-i, Extended Data Fig. 5). Given that PAX7 is expressed in a subset of neural crest cells in chicken limbs, we verified the absence of neural crest cell contribution to the muscle lineage using isotopic/isochronic GFP+chicken-into-chicken grafts of neural tubes and genetic lineage tracing experiments in limbs of *Wnt1*^Cre^:*R26*^*stop*^/^Tom^ mice^29^. Using these approaches, we did not find any contribution of neural crest cells to either myogenic cells or myonuclei, at muscle tips close to tendons in chicken and mouse limbs (Extended Data Fig. 6). The cell lineage tracing experiments in avians and mice led us to conclude that a fraction of myogenic cells at muscle tips close to tendons are derived from limb CT fibroblasts and are not somite/*Pax3*-derived.

Our results showing a recruitment of CT fibroblasts into the muscle lineage led us to reconsider the mechanisms underlying limb muscle patterning. We propose a fibroblast-to-myoblast conversion model, whereby a population of CT fibroblast progenitors acquire progressively a myogenic signature and are incorporated at the tips of existing myotubes during development. In this model, progenitors would exhibit a dual CT/muscle identity. To search for such cells with dual identity, we performed single-cell RNA-sequencing (scRNAseq) of chicken whole-limb cells at successive developmental stages: E4 a progenitor stage, E6 when major spatial re-arrangements occur for muscle and CTs and E10 when the final muscle pattern is set (Fig. 3). At each stage, we detected muscle and CT clusters that represented the majority of limb cells (90 to 95% of limb cells), in addition to other clusters encompassing the expected cell populations present in developing limb tissues such as vessels, blood, innervation, and ectoderm (Fig. 3a). Skeletal muscle clusters (which only include mononucleated cells) started from 2% at E4 and reached 29% of limb cells at E10, while CT clusters started from 96% at E4 and comprised 63% of limb cells at E10 (Fig. 3a). In addition to the CT and muscle clusters, we identified a subpopulation of cells co-expressing at least one CT marker, *PRRX1*^30^, *PDGFRA^31^, TWIST2^32^, OSR1^9^, SCX^24^* with one muscle marker, *PAX7, MYF5, MYOD, MYOG* (Fig. 3b, Extended Data Fig. 7a,b). This cell population exhibits a dual CT/muscle (CT/M) identity expressing the Top10 markers identified for CT and for muscle clusters (Fig. 3b). This dual-identity population increased from E4 (0.5% of limb cells) to reach 4% of limb cells at E6, when the major spatial re-arrangements occur for muscle and CTs, and was maintained at 3.3% at E10 when the final muscle and CT pattern is set (Fig. 3b). Among CT markers, *PRRX1* was expressed in the majority of CT/M cells at all stages (85% of CT/M cells at E4, 72% at E6 and 66% at E10). The fraction of *PRRX1*+ cells among CT/M cells was maintained, while the fraction of *PRRX1+* cells among all limb cells dropped with time from 88% at E4 to 25% at E10 (Extended Data Fig. 7c), suggesting that *PRRX1* expression was actively retained in CT/M cells. *TWIST2*, expressed in 15 to 29% of CT/M cells was the second major contributor to CT/M identity (Extended Data Fig. 7c). In contrast, the contribution of *SCX* to the CT/M identity decreased with time from 27% to 6% of CT/M cells (Extended Data Fig. 7c). Twenty to 30% of the CT/M cells expressed more than one CT marker and the major combination at all stages was *PRRX1/TWIST2*. Altogether, these results point to a key role for *PRRX1* and *TWIST2* in the CT/M identity and that a TWIST2 cell population contributes to developmental muscles in addition to adult skeletal muscles^32^. The myogenic markers, *PAX7* and *MYOD1* showed similar distributions in CT/M cells, as a majority of CT/M cells expressed PAX7 (42 to 57% of CT/M cells) and MYOD1 (36 to 57% of CT/M cells), while much fewer cells expressed MYOG (10 to 15% of CT/M cells). These observations are suggestive of an immature state of these bi-potent CT/M cells (Extended Data Fig. 7c).

**Figure 2.**
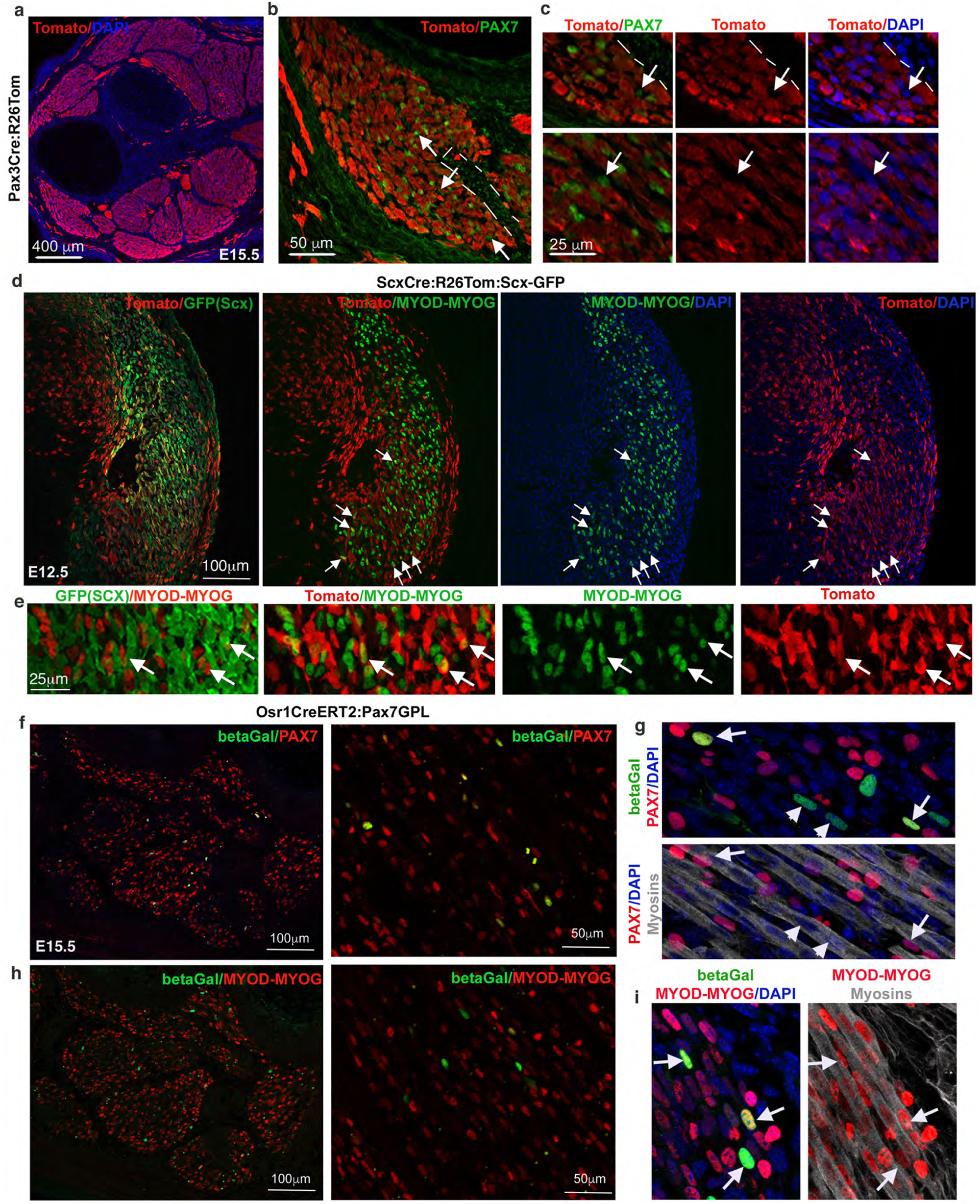
A subpopulation of muscle cells are not derived from the *Pax3* lineage but are derived from the fibroblast lineage in mouse limbs. (**a**) Transverse forelimb sections of E15.5 *Pax3^Cre^:R26^stop/Tom^* mice immunostained with Tomato antibody and DAPI staining. (**b**) Focus on a muscle immunostained with Tomato and PAX7 antibodies. (**c**) High magnifications showing PAX7+ cells that are tomato-negative i.e. not of *Pax3* lineage (arrows). (**d**) Transverse forelimb sections of E12.5 *Scx^Cre^:R26^stop/Tom^*:GFP(Scx) mice immunostained with Tomato (red) and MYOD-MYOG (green) antibodies, and stained with DAPI (blue). Combinations of two colours are indicated on panels. Arrows point to MYOD-MYOG+ cells that are Tomato+ (*Scx* lineage). (**e**) High magnifications showing MYOD-MYOG+ cells that are Tomato+ (*Scx* lineage). (**f,h**) Transverse and adjacent forelimb sections of E15.5 *Osr1^CreERT2^:Pax7^GPL^* mice immunostained with betaGal (*Osr1*-lineage-derived cells, green) and PAX7 (red) antibodies, combined with DAPI (blue) (f), and with betaGal (*Osr1*-lineage-derived cells, green) and MYOD/MYOG (red) antibodies combined with DAPI (h). (**g,i**) High magnifications. (g) Arrows point to the betaGal+ nuclei (*Osr1*-lineage-derived, green) and PAX7+ (red) that are located in between myotubes, while arrowheads point to the betaGal+ nuclei (green) that are PAX7- and located inside myotubes. These betaGal+ PAX7-nuclei correspond to cells in which PAX7 has been donwregulated and betaGal expression kept after cell incorporation into myotubes. (i) Arrows point to the betaGal+nuclei (*Osr1*-lineage-derived, green) that are MYOD/MYOG+ (red).

**Figure 3.**
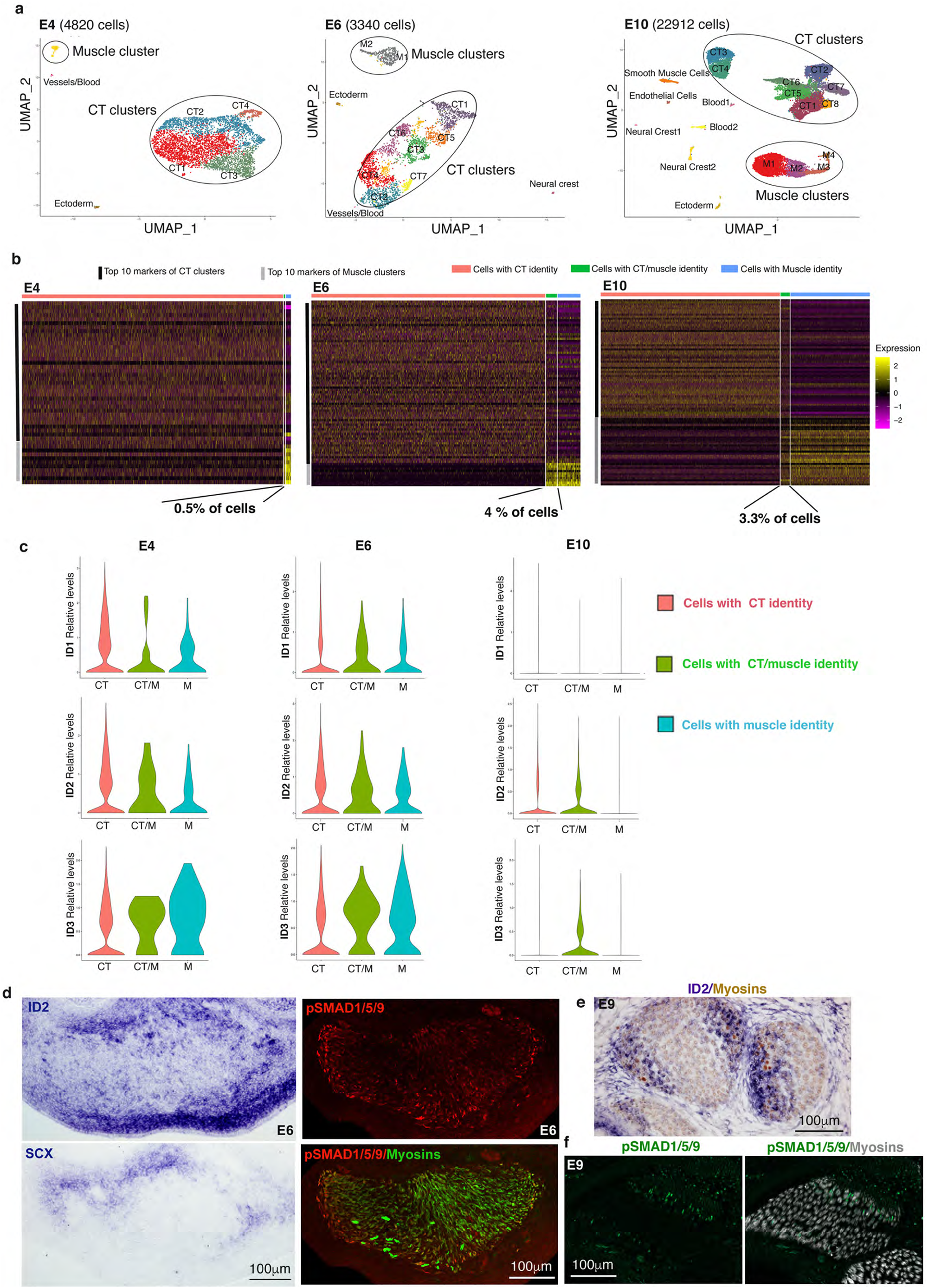
BMP signalling is active in cells displaying a CT/M transcriptional identity and in cells at the muscle/tendon interface in chicken limbs. (**a**) UMAP plots showing the whole-limb clustered populations with their respective number of cells at E4 (4820 cells), E6 (3340 cells) and E10 (22912 cells). CT and muscle clusters represent the majority of limb cells at each developmental stage. **(b)** Heatmaps showing the relative expression of the top 10 markers for each CT and muscle clusters in cells grouped by their identity CT (red), CT/M (green) or M (blue) at E4, E6 and E10. Upregulated genes in yellow, downregulated genes in purple. CT/M identity labels 26 cells (0.5% of limb cells) at E4, 133 cells (4.0% of limb cells) at E6 and 729 cells (3.3% of limb cells) at E10. (**c**) Violin plots showing Log-normalized expression levels of *ID1*, *ID2* and *ID3* genes, which are transcriptional readouts of the BMP activity, in cells grouped by their identity CT (red), CT/M (green) or M (blue) at E4, E6 and E10. **(d)** Adjacent and transverse limb sections of E6 chicken embryos were hybridized with ID2 or SCX probes and immunostained with pSMAD1/5/9 and MF20 (myosins) antibodies. High magnifications of ventral limb muscle masses. (**e**) Transverse limb sections of E9 chicken embryos were hybridized with ID2 probe (blue) and then immunostained with MF20 (myosins) antibody (brown). Focus on dorsal limb muscles. (**f**) Transverse limb sections of E9 chicken embryos were immunostained with pSMAD1/5/9 (green) and MF20 (myosins, grey) antibodies. Focus on dorsal limb muscles.

*ID* genes are recognized transcriptional readouts of BMP activity^33^. *ID2* and *ID3* genes were expressed in a similar or greater fraction of CT/M cells compared to CT and M cells (Fig. 3c). *ID1* showed an interesting expression kinetic as it peaks at E6 when muscle and CT spatial re-arrangements occur (Fig. 3c). Consistent with *ID* gene expression in this population with a dual identity (Fig. 3c), BMP activity visualized with *ID2* and pSMAD1/5/9 expression was enriched at the CT/muscle interface, in between muscle masses and tendon primordia at E6 (Fig. 3d) and at muscle tips close to tendon at E8/E9 (Fig. 3e,f, Extended Data Fig. 8). Interestingly, we detected 2 populations of pSMAD1/5/9+ myonuclei at muscle tips, one somite-derived (yellow arrows in Extended Data Fig. 9) and one non-somite-derived (white arrows in Extended Data Fig. 9) at both E6 and E9 stages. This is reminiscent of the two distinct myonuclear populations recently identified in adult mouse muscle at the myotendinous junction, with one being enriched with CT-associated collagen genes^34^; leading to the compelling notion that the myonuclei with a fibroblast signature could be of CT origin. Altogether, these results show the existence of a BMP-responsive population with a dual CT/muscle identity that could correspond to the fibroblast-derived myogenic cells found at muscle tips close to tendon.

Based on BMP activity at the CT/muscle interface (Fig. 3d-f, Extended Data Fig. 8) and in a cell subpopulation with a dual CT/M identity (Fig. 3c) in chicken limbs, we hypothesised that BMP signalling would be involved in the recruitment of a subpopulation of CT fibroblasts towards a muscle fate. To test the ability of BMP signalling to control the fibroblast-to-myoblast conversion, we overexpressed BMP activity in chicken limbs and assessed the consequences for PAX7+ myogenic progenitors and TCF4+ CT fibroblasts (Fig. 4a-g). Exposure of chicken limbs to retroviral BMP4 led to an increase in the number of PAX7+ muscle progenitors (Fig. 4a,c,f) as previously observed^21^. However, consistent with the cell fate conversion hypothesis, the increase of PAX7+ progenitors occurred at the expense of TCF4+ fibroblasts (Fig. 4a-f). The inverse correlation between PAX7+ and TCF4+ cells upon BMP exposure was not accompanied by changes in the total cell number (Fig. 4g). Moreover, no change in the proliferation rate of PAX7+ cells was observed upon BMP4 exposure in chicken limbs (Extended Data Fig. 10a-g). This abnormal increase in PAX7+ cells led to muscle patterning defects as BMP4 overexpression induced muscle splitting defects, visualised by muscle fusion (Fig. 4a-c,e,f), due to the reduction of TCF4+ fibroblasts known to control muscle patterning^17^. Because there is no patterning process *in vitro*, BMP overexpression was then applied to myoblast primary cultures in proliferation culture conditions. Consistent with our conversion hypothesis and patterning defects observed *in vivo*, BMP4 treatment did not alter the number of PAX7+ cells in myoblast cultures (Extended Data Fig. 10h-j). Our interpretation is that, in contrast to the *in vivo* situation where BMP-competent fibroblasts are present to become myoblasts upon BMP exposure, the contaminating fibroblasts inherent to primary myoblast cultures were not present in enough quantity to generate a detectable conversion phenotype. The absence of any BMP4 effect on the number PAX7+ cells in myoblast cultures is in support of the fibroblast-to-myoblast conversion upon BMP4 exposure in chicken limbs. To mimic the *in vivo* conversion, we applied retroviral BMP4 or BMPR1Aca (a constitutive active form of the BMPR1A) to CEFs (chicken embryonic fibroblasts) that were isolated from whole embryos. BMP over-activation induced the appearance of PAX7+ cells compared to control cultures, albeit with a low efficacy (Fig. 4h,i). This indicates that a subset of this mixed fibroblast population can convert into PAX7+ cells when exposed to BMP *in vitro*.

**Figure 4.**
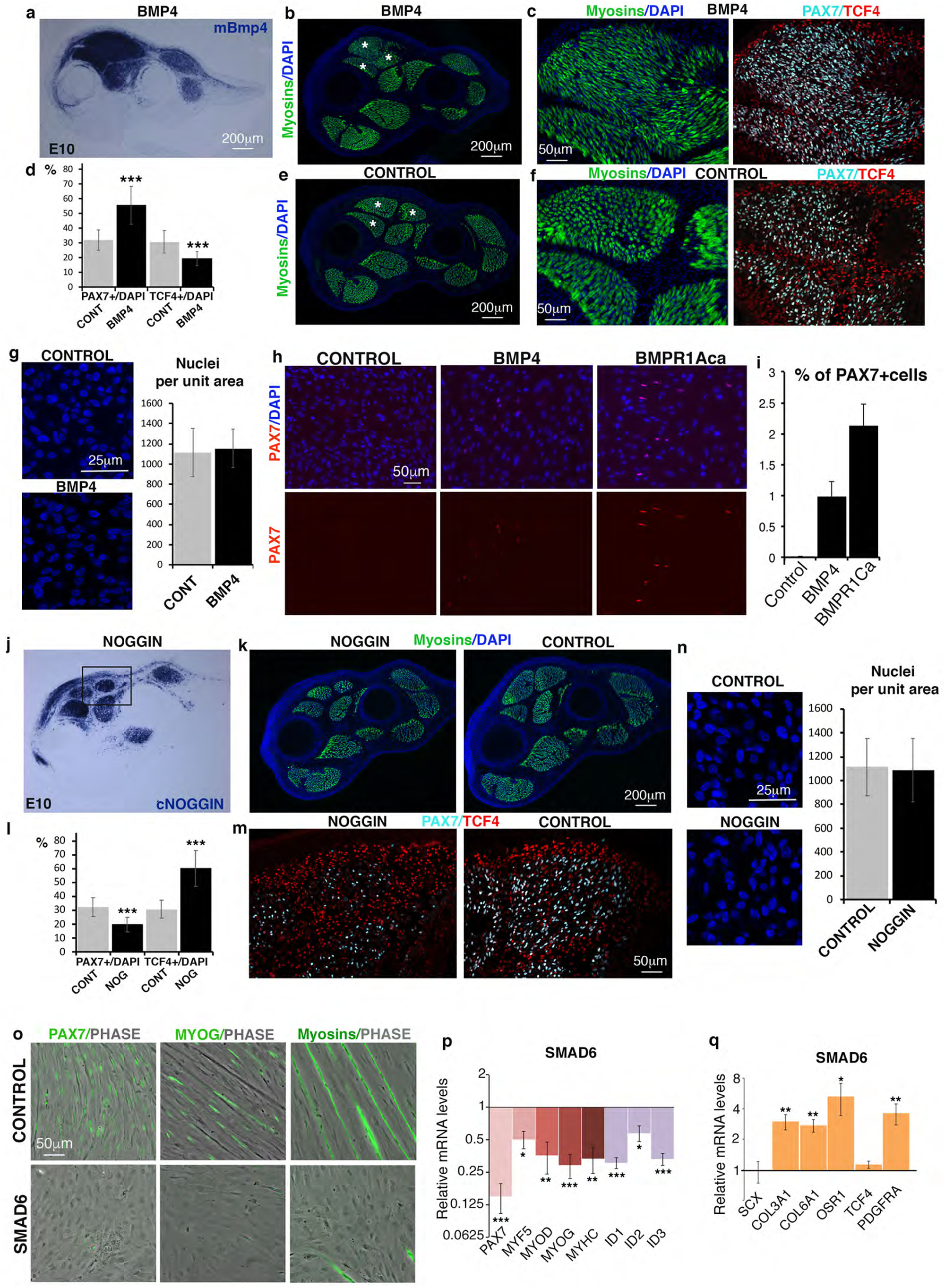
BMP signalling regulates a fibroblast-myoblast conversion in vivo and in vitro. **(a-g)** BMP4/RCAS-producing cells were grafted to right limbs of E5 chicken embryos. Embryos were fixed 5 days later at E10. *In situ* hybridisation experiments with mBmp4 probes to transverse limb sections indicate the extent of ectopic BMP4 in dorsal limb muscles (**a**). (**b,e**) Limb sections were immunostained with MF20 (myosins, green) to visualise the muscle pattern. Stars indicate fused muscles in BMP4-treated limbs (b) versus individual muscles in control limbs (e). (**c,f**) High magnifications of dorsal limb muscles immunostained with MF20 (myosins, green), PAX7 (cyan) and TCF4 (red) antibodies in BMP4 (c) and respective control limbs (f). (**d**) Percentages of PAX7+ and TCF4+ nuclei per total number of nuclei in BMP4-treated limbs versus controls. (**g**) Representative fields of DAPI+ nuclei in muscles of control and BMP4 limare and quantification of the number of nuclei per unit area. **(h**) Chicken fibroblast cultures treated with BMP4/RCAS or BMPR1ca/RCAS. Immunostaining with PAX7 antibody (red) with DAPI in BMP4- and BMPR1ca-treated fibroblast cultures compared to controls. (**i**) Quantification of PAX7+ cells versus the total cell number. (**j-n**) NOGGIN/RCAS-producing cells were grafted to right limbs of E5 chicken embryos. Embryos were fixed 5 days later at E10. (**j**) *In situ* hybridisation experiments with cNOGGIN probes to transverse limb sections indicate the extent of ectopic NOGGIN in dorsal limb muscles. (**k**) Limb sections were immunostained with MF20 (myosins, green) to visualise muscles. (**m**) High magnifications of dorsal limb muscles immunostained with PAX7 (cyan) and TCF4 (red) antibodies in NOGGIN and respective control limbs. (**l**) Percentages of PAX7+ and TCF4+ nuclei per total number of nuclei in NOGGIN-treated limbs versus controls. (**n**) Representative fields of DAPI+nuclei in muscles of control and NOGGIN limbs; and quantification of the number of nuclei per unit area. (**o-q**) Myoblast cultures treated with SMAD6/RCAS were induced to differentiation. (**o**) Immunostaining with PAX7, MYOG, and MF20 (myosins) antibodies in control and SMAD6 cultures. (**p,q**) RT-qPCR analyses of the expression levels of muscle genes, *PAX7, MYF5, MYOD, MYOG, MYHC*, BMP-responsive genes *ID1, ID2, ID3* (p) and connective tissue genes, *SCX, COL3A1, COL6A1, OSR1, TCF4, PDGFRA* (q) in SMAD6-treated and control cell cultures. Gene mRNA levels were normalized to *GAPDH* and *RPS17*. The relative mRNA levels were calculated using the 2^^−ΔΔCt^ method using the control condition as controls. For each gene, the mRNA levels of the control condition were normalized to 1. Graph shows means ± standard deviations of 9 biological samples (p) and 6 biological samples (q).

BMP loss-of-function experiments in chicken limbs, by overexpression of the antagonist NOGGIN induced the converse phenotype, *i.e*. an increase in the number of TCF4+ fibroblasts at the expense of PAX7+muscle progenitors with no change in cell number (Fig. 4j-n). The behaviour of TCF4+ fibroblasts upon BMP misexpression is consistent with the decrease and increase in the expression of the *OSR1* CT marker upon BMP gain- and loss-of-function experiments in chicken limbs, respectively^35^. Consistent with the *in vivo* experiments in chicken embryos, the inhibition of BMP signalling in myoblast cultures led to a fibroblast phenotype (Fig. 4o-q, Extended Data Fig. 10k). BMP inhibition was achieved with the overexpression of SMAD6 (Fig. 4o-q), an inhibitor of the pSMAD1/5/8 pathway^33^ or the BMP antagonist NOGGIN (Extended Data Fig. 10k). BMP inhibition led to a reduction of PAX7+, MYOD+ and MYOG+ cells and myotubes in myoblast cultures in differentiation culture conditions (Fig. 4o). This was associated with a decrease in the mRNA expression levels of muscle lineage markers from progenitor to differentiation state (*PAX7, MYF5, MYOD, MYOG, MYHC*) (Fig. 4p). The SMAD6-treated muscle cells adopted a fibroblast-like phenotype compared with myotubes observed in control myoblast cultures (Fig. 4o) and displayed an increased expression of CT markers, such as *COL3A1, COL6A1* and *OSR1* (Fig. 4q). These results demonstrate that BMP inhibition converts myoblasts towards a fibroblastic phenotype.

Altogether, these *in vivo* and *in vitro* BMP functional experiments unravel an unexpected role for BMP signalling in driving a fibroblast/myoblast switch during limb muscle development.

### Conclusion

This work shows that the tendon/muscle interface partly results from a fibroblast-to-myoblast conversion driven by localized BMP signalling. These data lead us to propose a scenario in which BMP locally drives the conversion of CT fibroblasts to myoblasts that are incorporated into myotubes at muscle tips close to tendons. This process leads to myonuclei of fibroblast origin at the tendon/muscle interface. Persistent BMP signalling regionalised at muscle tips then provides an *in vivo* safeguard to maintain the muscle identity of fibroblast-derived muscle cells. Previous BrdU/EdU-based experiments have shown a preferential incorporation of nuclei in myotubes at muscle tips during foetal and postnatal myogenesis^36–38^. Although these BrdU/EdU-based experiments never addressed the origin of the newly incorporated nuclei, it was always assumed that this preferential incorporation at the muscle tips was with somitic-derived nuclei. We now propose that the preferential nucleus incorporation at muscle tips of myotubes occurs with nuclei of fibroblast origin. We see these muscle cells of CT fibroblast origin are a source of positional information for muscle patterning, possibly regulating the localisation of myoblast fusion events and thereby the shaping and orientation of growth of limb skeletal muscles during foetal development. These muscle cells of fibroblast origin are reminiscent of muscle founder cells that drive muscle identity in *Drosophila* embryos^39^. Future studies will explore whether this mechanism can be reactivated in adult where it could be relevant in the context of muscle regeneration.

## Material and methods

### Chicken and quail embryos

Fertilized chicken and quail eggs from commercial sources (White Leghorn and JA57 strain, Japanese quail strain, Morizeau, Dangers, France) were incubated at 37.5°C. Before 2 days of incubation, chicken and quail embryos were staged according to somite number. After 2 days of incubation, embryos were staged according to days of *in ovo* development. GFP+chicken were generated and provided by the Roslin Institute^40^.

### Mouse strains

Animals were handled according to European Community guidelines and protocols were validated by the ethic committee of the French Ministry. Embryos were dated taking the day of the vaginal plug as E0.5. Mouse lines used in this study were kept on a mixed C57BL/6JRj and DBA/2JRj genetic background (B6D2F1, Janvier862 Labs). The following strains were described previously: *Pax3^Cre^:R26^Tomato^* (Ai9)^26,41^, *Wnt1*^*Cre*29^, *Scx*^*Cre*42^, *Osr1^GCE^* allele (*Osr1^eGFP-CreERT2^*)^28^, *Pax7^GPL^* reporter (*Pax7^nGFP-stop-nlacZ^*)^27^ and *Scx^Cre^:R26^Tomato^* (Extended Ressources Table 1). For temporal fate mapping using the *Osr1^GCE^* allele, we used the lineage-specific Pax7-driven reporter (*Pax7^GPL^*), in which locus accessibility to Cre-mediated recombination allows higher sensitivity for tracking Pax7-expressing cells and their progeny^27^. To induce recombination, 5 mg of tamoxifen (Sigma #T5648) were administered by gavage to pregnant females. A 25 mg/ml stock solution in 5% ethanol and 95% sunflower seed oil was prepared by thorough resuspension with rocking at 4°C.

### Surgical grafting experiments in avian embryos

#### Isotopic/Isochronic quail- or GFP+chicken-into-chicken grafts of presomitic mesoderm

Quail, GFP+chicken and chicken embryos were incubated until they reached 15-somite stage. Of note, limb buds are not formed at this stage and brachial somitic cells have not yet migrated into the forelimb presumptive regions. An artificial dark field was obtained by injecting Indian ink, diluted 1:1 in PBS, beneath the chicken host embryos. Microsurgery was performed on the right side of the host embryos. The 15^th^ somite and the non-segmented presomitic mesoderm was removed over a length corresponding to 5–8 somites. Presomitic mesoderms from either quail (N=6) or GFP+chicken (N=4) donor embryos were rinsed in DMEM (PAA Laboratories)/10% foetal calf serum (FCS, Eurobio) and transplanted into chicken hosts submitted to the same ablation. Grafts were performed according to the original dorso-ventral and antero–posterior orientations (Fig. 1a-g, Extended Data Fig.1,2,9).

#### Isotopic/Isochronic quail-into-chicken grafts of limb lateral plate

Quail and chicken embryos were incubated until they reached 18-somite stage. Of note, limb buds are not formed at this stage, and brachial somitic cells have not yet migrated intro the forelimb lateral plate. The forelimb lateral plate mesoderm at the level of the 15^th^ somite and over a length of 3 somites was excised from host chicken embryos and replaced by their quail counterpart (N=3) (Fig. 1h-k, Extended Data Fig.3).

#### Isotopic/isochronic GFP+chicken-into-chicken grafts of neural tube

GFP+chicken and chicken embryos were incubated until they reached 12- to 14-somite stage. At this stage, neural crest cells did not yet leave the neural tube to colonize the limb bub. The neural tube was excised over a length corresponding to 5–8 somites from GFP+ chicken donor embryos and grafted in place of the neural tube of chicken host embryos (N=4) (Extended Data Fig. 6).

For all types of grafts, quail-chicken and GFP+chicken-chicken chimeras were allowed to grow for another 4 to 8 days and treated for either immunohistochemistry or *in situ* hybridization to tissue sections.

### Grafts of BMP4-RCAS and NOGGIN-RCAS expressing cells in chicken limbs

Chicken embryonic fibroblasts (CEFs) obtained from E10 chicken embryos were transfected with BMP4-RCAS or NOGGIN-RCAS using the Calcium Phosphate Transfection Kit (Invitrogen, France). Cell pellets of approximately 50–100 μm in diameter were grafted into the limbs of E5 embryos as previously described^21^. BMP4-RCAS- (N=8) and NOGGIN-RCAS- (N=4) grafted embryos were harvested at E9/E10 and processed for *in situ* hybridization and immunohistochemistry. Cell counting was performed with Fiji software.

### Immunohistochemistry

For antibody staining, control and manipulated chicken forelimbs were fixed in a paraformaldehyde 4% solution, embedded in gelatin/sucrose and then cut in 12-μm cryostat sections. All antibodies (sources, conditions of use, references) are reported in Extended Ressources Table 1. Quail cells were detected using the QCPN antibody. Differentiated muscle cells were detected using the monoclonal antibody, MF20 recognizing sarcomeric myosin heavy chains. Myoblasts were detected using the MYOD and MYOG antibodies. Muscle progenitors were detected using the monoclonal PAX7 antibody. Active BMP signalling was detected using the polyclonal pSMAD antibody recognizing the complex BMP-activated receptor-phosphorylated SMAD1/5/9. Tendons were detected using Collagen type XII antibody. The following secondary antibodies (Molecular probes) were used: goat anti-mouse IgG coupled to Alexa fluor 555, goat anti-mouse IgG1 coupled to Alexa fluor 568, goat anti-mouse IgG2 coupled to Alexa fluor 488, and goat anti-rabbit IgG coupled with Alexa fluor 488 or 555. To label nuclei, sections were incubated for 15 minutes with DAPI. Stained sections were examined using the Apotome.2 microscope (Zeiss) or LSM700 confocal microscope (Zeiss).

### *In situ* hybridization to tissue sections

Normal or manipulated embryos were fixed in a paraformaldehyde 4% solution. Limbs were cut in 12-μm cryostat transverse sections and processed for *in situ* hybridization. Alternating serial sections from embryos were hybridized with probe 1 and probe 2. The digoxigenin-labelled mRNA probes were used as described: SCX, ID2, BMP4, NOGGIN (chicken probes), and mBmp4 (mouse probe)^21,35^.

### Single-cell RNA-sequencing analysis from limb cells

For scRNAseq analysis of chicken limb cells at different developmental stages, forelimbs from 2 different E4 embryos, 3 different E6 embryos and 3 different E10 embryos were dissected and dissociated by collagenase and mechanical treatments. Cell concentration was adjusted to 5000 cells/μl in buffer. 5000 cells per conditions were loaded into the 10X Chromium Chip with the Single Cell 3’ Reagent Kit v3 according to the manufacturer’s protocol. Samples were pooled together at each developmental stage (E4 (N=2), E6 (N=3) and E10 (N=3)) before making the libraries. Libraries were then sequenced by pair with a HighOutput flowcel using an Illumina Nextseq 500 with the following mode (150 HO): 28 base-pairs (bp) (Read1), 125 bp (Read 2) and 8 bp (i7 Index). A minimum of 50 000 reads per cell were sequenced and analyzed with Cell Ranger Single Cell Software Suite 3.0.2 by 10X Genomics. Raw base call files from the Nextseq 500 were demultiplexed with the cellranger mkfastq pipeline into library specific FASTQ files. The FASTQ files for each library were then processed independently with the cellranger count pipeline. This pipeline used STAR21 to align cDNA reads to the *Gallus gallus* genome (Sequence: GRCg6a, *Gallus gallus* reference). The used sample-size in scRNAseq allowed us to robustly identify cell populations as small as 18 cells at E4, 6 cells at E6 and 66 cells at E10.

The Seurat package (v3.0) was used to perform downstream clustering analysis on scRNAseq data^43^. Cells went through a classical Quality Control using the number of detected genes per cell (nFeatures), the number of mRNA molecules per cell (nCounts) and the percentage of expression of mitochondrial genes (pMito) as cut-offs. Outliers on a nFeature vs nCount plot were manually identified and removed from the dataset. Most importantly for this study, potential doublets were identified by running the Scrublet algorithm^44^ and then removed from the dataset. Gene counts for cells that passed the above selections were normalized to the total expression and Log-transformed with the NormalizeData function of Seurat using the nCount median as scale factor. Highly variable genes were detected with the FindVariableFeatures function (default parameters). Cell cycle effect was regressed out using the ScaleData function. Using highly variable genes as input, principal component analysis was performed on the scaled data in order to reduce dimensionality. Statistically significant principal components were determined by using the JackStrawPlot and the ElbowPlot functions. Cell clusters were generated with the FindNeighbors/FindClusters functions (default parameters except for the number of selected PCs). Different clustering results were generated at different resolutions and for different sets of PCs. Non-linear dimensional reduction (UMAP) and clustering trees using Clustree (Zappia 2018 Gigascience doi:gigascience/diy083) were used to visualize clustering results and select for the most robust and relevant result. Differentially expressed genes were found using the FindAllMarkers function of Seurat (using highly variable genes as an input, default parameters otherwise) that ran Wilcoxon rank sum tests.

For the analysis of the CT/M cells, cells were grouped according to 4 identities (CT, CT/M, M and Other). The CT/M identity is defined by the co-expression (i.e gene Log-normalized count > 0) within a cell of at least one of the CT markers (PRRX1, TWIST2, PDGFRA, OSR1, SCX) with at least one of muscle markers (PAX7, MYF5, MYOD1, MYOG). CT identity is conferred to all cells allocated to a CT cluster, except the CT/M cells. M identity is conferred to all cells allocated to a muscle cluster, except the CT/M cells. « Other » identity is conferred to all cells to clusters that are neither CT nor muscle, except for the CT/M cells. The scRNAseq datasets were then analysed with the angle of these 4 identities using Seurat tools such as Feature plots, Heatmaps and Violin plots.

### Cell cultures

Chicken embryonic fibroblast (CEF) cultures were obtained from E10 chicken embryos. Briefly, 10 embryos without heads and viscera were mechanically dissociated. After centrifugation, cells were plated. Myoblast primary cultures were obtained from limbs of E10 chicken embryos as already described^45^.

Empty/RCAS, BMP4/RCAS, BMPR1ACa/RCAS^21^ were transfected to primary fibroblast cultures. Transfected fibroblasts were left for 5 to 7 days in culture with cell splitting to allow virus spread. Empty/RCAS, BMP4/RCAS, BMPR1ACa/RCAS fibroblasts were then fixed and processed for immunohistochemistry.

Empty/RCAS, BMP4/RCAS, NOGGIN/RCAS^21^ and SMAD6/RCAS were transfected to myoblast primary cultures. Transfected myoblasts were left for 5 to 7 days in culture with cell splitting to allow virus spread. Transfected myoblasts were either analysed in proliferation or in differentiation culture conditions. Empty/RCAS, NOGGIN/RCAS, SMAD6/RCAS myoblasts were fixed and processed for immunohistochemistry or RT-q-PCR assays to analyse gene expression.

### Quantitative real-time PCR

qPCR was performed as described in^22^. Total RNAs were extracted from SMAD6 or control myoblasts. 500 ng of RNA was reverse-transcribed. qPCR was performed using SYBR Green PCR master Mix (Aplied Biosystems) with primers listed in Extended Ressources Table 2. The relative mRNA levels were calculated using the 2^-ΔΔCt method^46^. The ΔCt were obtained from Ct normalized with GAPDGH and RPS17 levels in each sample.

## Statistics

GraphPad Prism 6 software was used for statistical analysis. Non-parametric tests were used to determine statistical significance, which was set at p values <0.05.

## GEO data accession number

Raw sequencing data will be deposited in the NCBI Gene Expression Omnibus database (https://www.ncbi.nlm.nih.gov/geo/) upon acceptance.

## Code availability

Data analysis was performed with standard pipelines in Seurat R packages, as described in the Methods. Scripts will be made available upon request.

## Acknowledgements

We thank Sophie Gournet for illustration. This work was supported by the CNRS, Inserm, SU, AFM and FRM. We thank the Roslin Intitute (Prof Helen Sang and Dr. Adrian Sherman) for providing us with GFP^+^ chicken eggs. The production of the GFP^+^ chicken embryos was supported by grants from BBSRC and the Wellcome Trust. ARTbio was supported by the CNRS, SU, the Institut Français de Bioinformatique (IFB) and by a grant from the SIRIC CURAMUS.

## Author contributions

JEL initiated the project, conducted the BMP experiments and contributed to Pax3 and Scx mouse work. CB conducted lateral plate grafting experiments, in situ hybridization and immunohistochemistry to experimental chicken and mouse limbs. MAB contributed to somite grafting experiments and performed cell preparation for scRNAseq. EH performed confocal imaging of chicken embryos and conducted the bioinformatic analysis of the scRNAseq datasets. GC conducted the mouse work with the Osr1, Pax3, Wnt1 drivers and immunohistochemistry with MYOG/MYOD antibodies. LY, CR performed GFP chicken (somite and neural tube) and quail (somite) grafting experiments. LB, SM contributed to the design of the scRNAseq analysis methods. RS provided the Scx:Cre embryos. SN, CFT performed qPCR on fibroblasts. JEL, EH, GC, LY, CR, CFT, ST, FR discussed about the project and contributed to write the manuscript. DD supervised the study, analysed the results, pictured most of the data and organised the figures, wrote the manuscript and acquired funding.

## EXTENDED DATA

**Extended Data Fig. 1.**
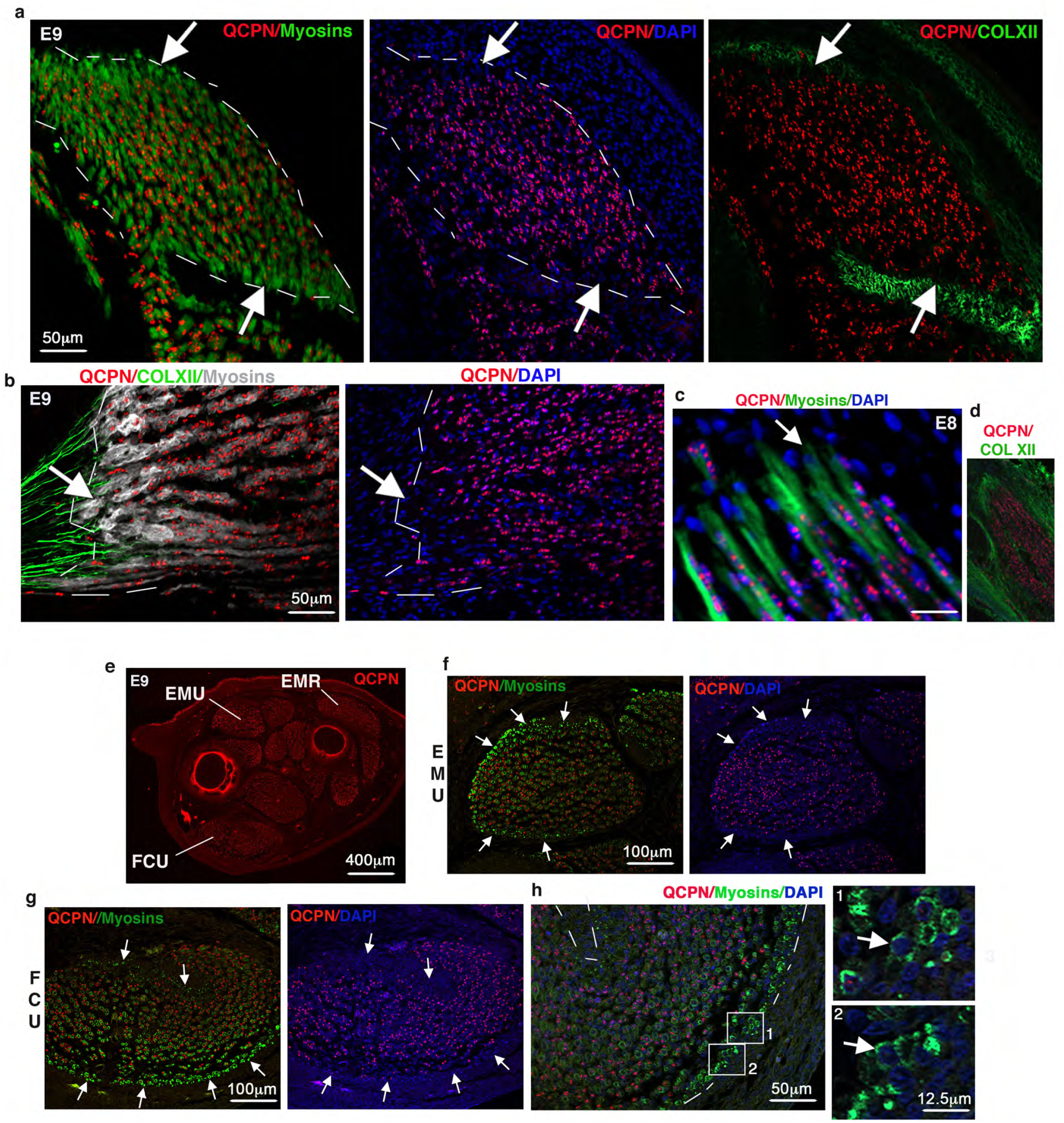
A subpopulation of myonuclei in skeletal muscles are not derived from presomitic mesoderm in chicken limbs. (**a,b**) Forelimbs of presomitic grafts of E9 quail-into-chicken embryos were co-immunostained with QCPN (quail nuclei), MF20 (myosins) and ColXII (tendons) antibodies. Focus on (a) the EMR (anterior muscle) and (b) a longitudinal view of muscle. Arrows point to zones of nuclei (DAPI) that are not of quail origin at the muscle tips close to tendons. (**c,d**) Focus on muscle tips from forelimb sections of presomitic grafts fixed at E8, immunostained with QCPN (quail nuclei), MF20 (myosins) and ColXII (tendons) antibodies. (**e-h**) Presomitic graft (quail-into-chicken embryos) fixed at E9. (**e**) Transverse forelimb sections immunostained with QCPN (quail nuclei). (**f**) Focus on the EMU. (**g,h**) Focus on the FCU. (h) Zoom to muscle tip regions with high magnifications showing quail-negative myonuclei (*i.e*. not presomitic-derived).

**Extended Data Fig. 2.**
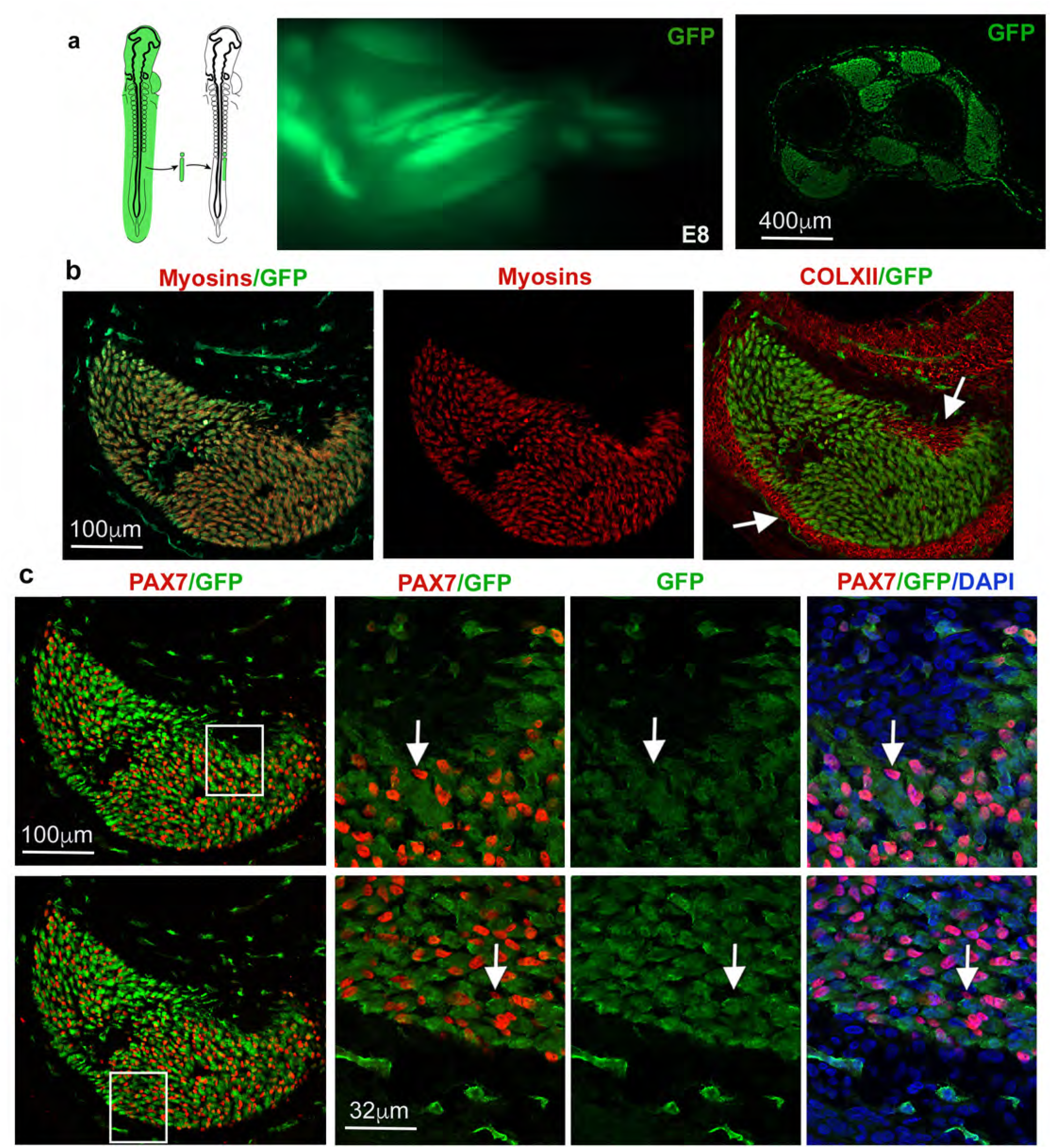
A subpopulation of PAX7+ cells are not derived from presomitic mesoderm in chicken limbs. (**a**) Diagram of presomitic grafts of GFP+chicken into chicken embryos (left panel). GFP fluorescence in forelimb of a presomitic graft E8 chimera (middle panel). Transverse limb section showing cytoplasmic GFP expressed in presomitic-derived cells in muscles (right panel). (**b**) Focus on FCU muscle labelled with GFP (green), MF20 (myosins, red) and ColXII (tendons, red) antibodies. Arrows point to tendons. (**c**) FCU muscle labelled with PAX7 (red) and GFP (green). High magnifications of FCU muscle tips show PAX7+ cells (arrows) that are GFP negative (*i.e*. not presomitic-derived).

**Extended Data Fig. 3.**
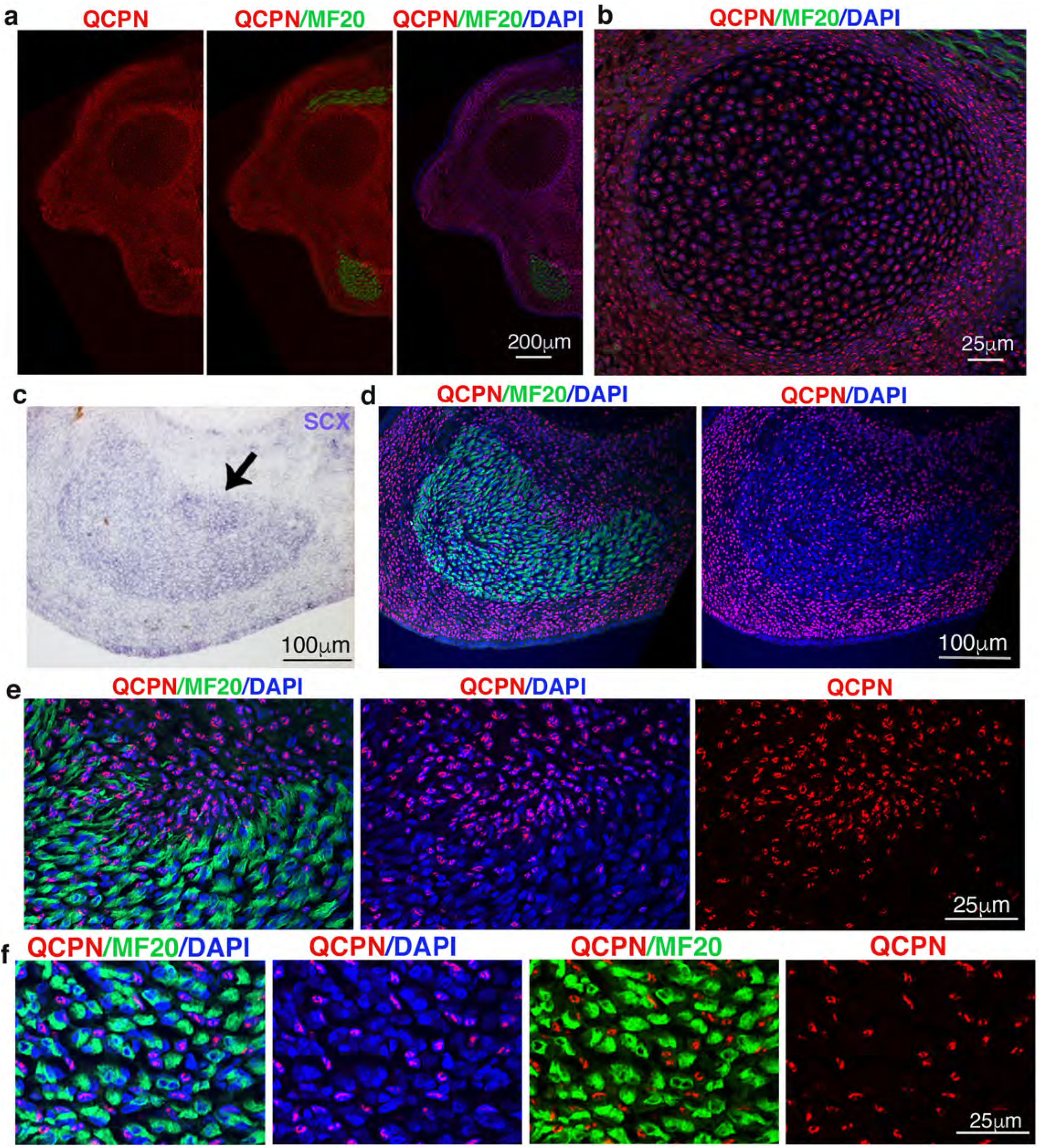
Quail lateral plate mesoderm grafts into chicken embryos lead to expected derivatives in limbs. (**a**) Limb transverse sections of lateral plate mesoderm grafts quail-into-chicken embryos fixed at E9 immunostained with QCPN (quail nuclei, red) and MF20 (myosins, green) antibodies, combined with DAPI. The quail+ nuclei (red), which are lateral plate-derived, are observed in cartilage (**b**), tendon (**d,e**), visualised with *SCX* expression by in situ hybridization on adjacent section (**c**) and in irregular CT in between myosin+ cells inside muscle (**f**). (e) is a zoom of the tendon regions of the FCU muscle (d).

**Extended Data Fig. 4.**
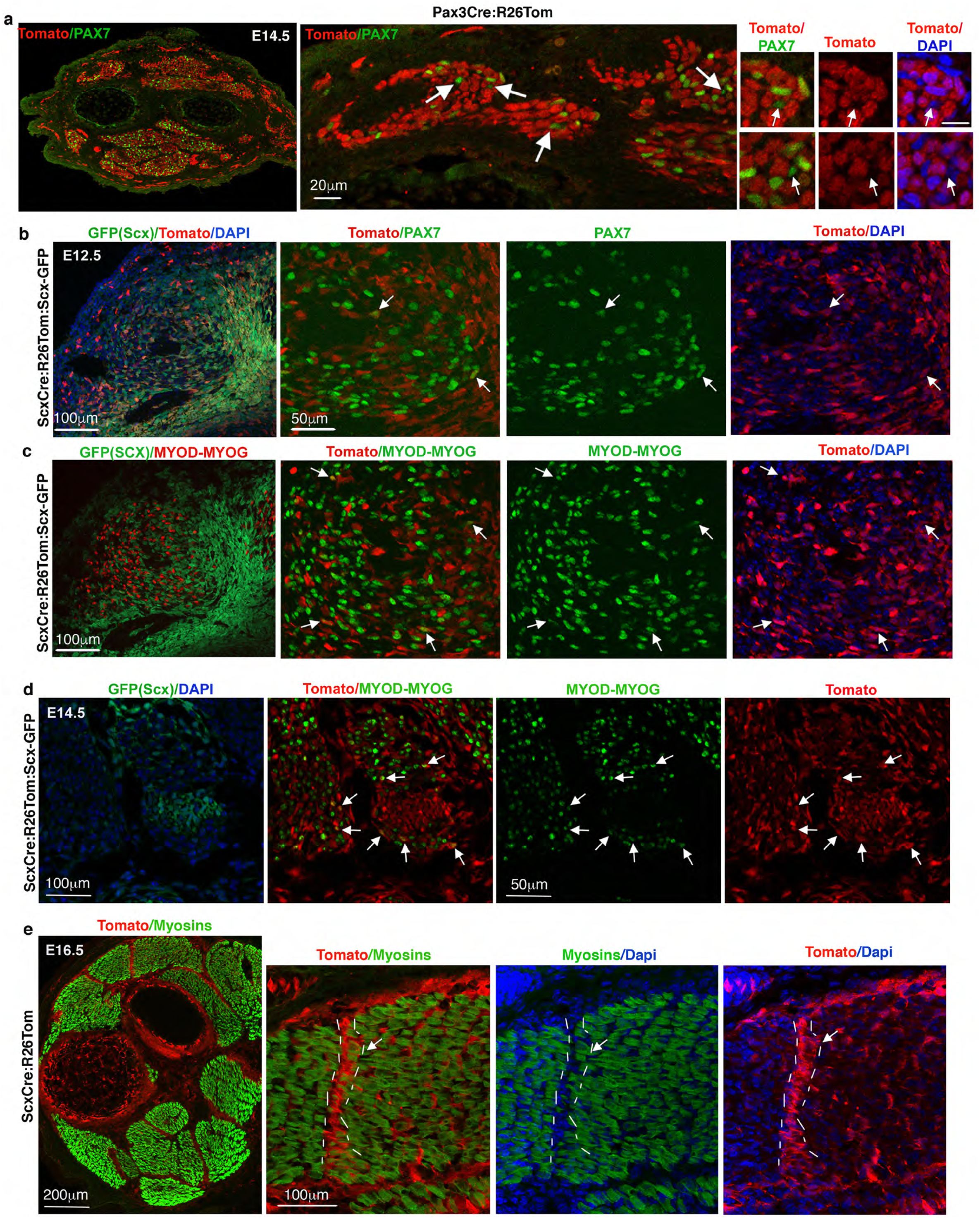
A subpopulation of PAX7+ progenitors and MYOD/MYOG+ myogenic cells are derived from *Scx*-lineage in mouse limbs. (**a**) Transverse forelimb sections of E14.5 *Pax3^Cre^:R26^Tom^* mice immunostained with Tomato (red) and PAX7 (green) antibodies, combined with DAPI staining. Arrows point to the PAX7+ cells (green) that are Tomato-negative (*i.e* not Pax3 lineage-derived). **(b)** Forelimb sections of E12.5 *Scx^Cre^:R26^Stop^/^Tom^:GFP(Scx)* mice immunostained with Tomato (Scx-lineage-derived cells, red) and PAX7 (green) antibodies. First panel shows GFP reflecting *SCX* expression in tendon primordia. Arrows point to Tomato+ cells (Scx-derived) that are PAX7+. **(c)** Adjacent sections of (b) immunostained with Tomato (Scx-lineage-derived cells, red) and MYOD/MYOG (green) antibodies. Arrows point to Tomato+ cells (Scx-derived) that are MYOD/MYOG+. First panel shows MYOD/MYOG+ (red) in between GFP+ cells (green). (**d**) Forelimb sections of E14.5 *Scx^Cre^:R26^Tom^:GFP(Scx)* mice were immunostained with Tomato (Scx-lineage-derived cells, red) and MYOD/MYOG (green) antibodies. Focus on ventral forelimb regions. First panel shows GFP reflecting *SCX* expression in tendons. Arrows point to Tomato+ cells (Scx-derived) that are MYOD/MYOG+ closed to tendons. **(e)** Transverse forelimb sections of E16.5 *Scx^Cre^:R26^Tom^* mice immunostained with Tomato (red) and My32 (myosins, green) antibodies. High magnifications of a Tomato+ junctional region between two dorsal limb muscles (encircled with dashed lines), showing the overlap between myosins and Tomato labelling (Scx-lineage-derived cells).

**Extended Data Fig. 5.**
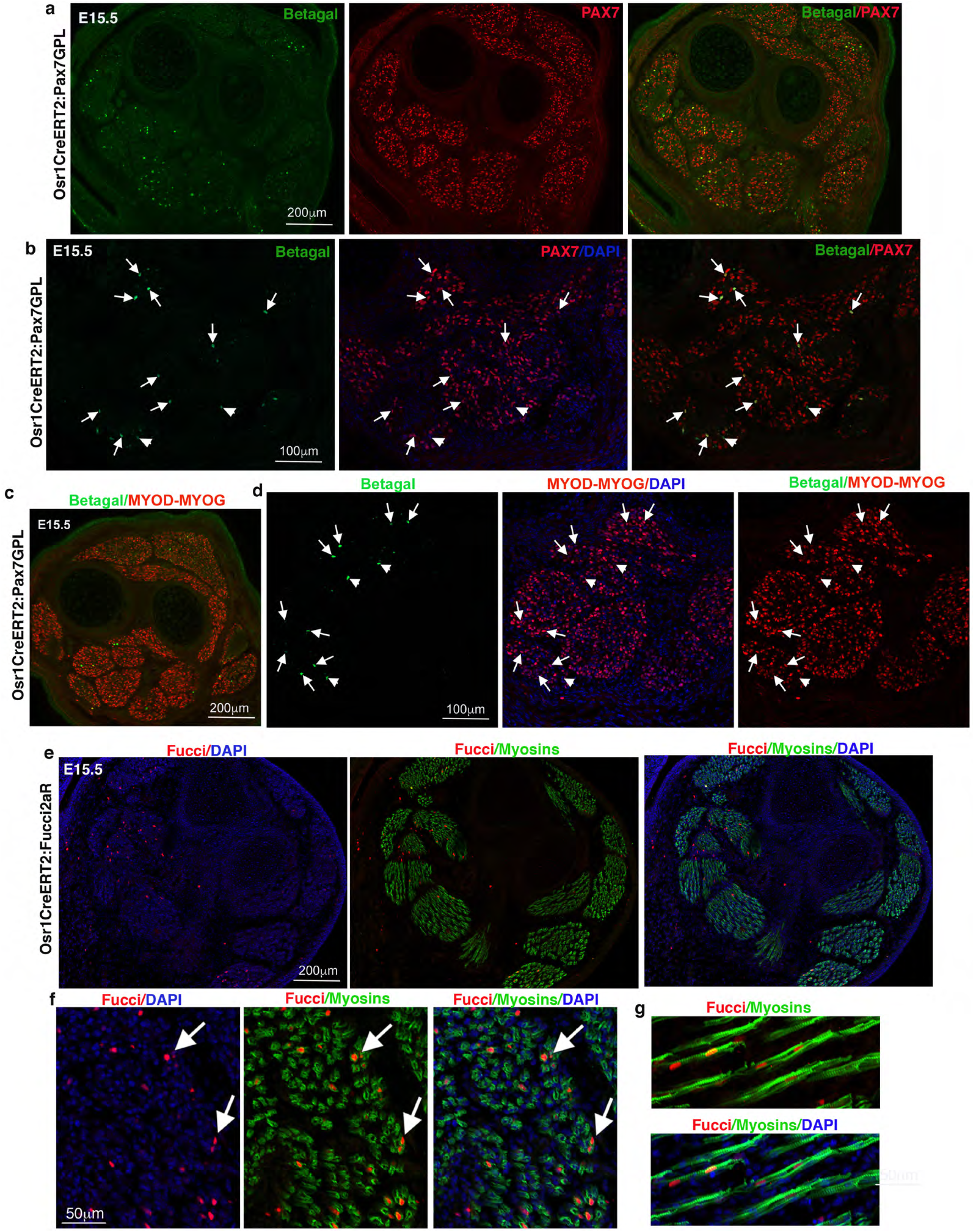
A fraction of muscle progenitors, myogenic cells and myonuclei are derived from *Osr1*- lineage in mouse limbs. (**a**) Transverse forelimb sections of E15.5 *Osr1^CreERT2^:Pax7^GPL^* mice immunostained with betaGal (Osr1-lineage-derived cells) and PAX7 antibodies. (**b**) High magnifications of ventral limb muscles. Arrows point to the betaGal+nuclei (green) that are PAX7+ (red), while arrowheads point to the betaGal+nuclei (green) that are not PAX7+. (**c-d**) Transverse forelimb sections of E15.5 *Osr1^CreERT2^:Pax7^GPL^* mice immunostained with betaGal (Osr1-lineage-derived cells) and MYOD/MYOG antibodies. (d) High magnifications of ventral limb muscles. Arrows point to the betaGal+nuclei (green) that are MYOD/MYOG+ (red), while arrowheads point to the betaGal+nuclei (green) that are not MYOD/MYOG+. (**e-f**) Transverse forelimb sections of E15.5 *Osr1^CreERT2^:Fucci2aR* mice immunostained with Fucci (Osr1-lineage-derived cells) and My32 (myosins) antibodies. (f) Arrows point to Fucci+nuclei (red) in myosin+ cells (green).

**Extended Data Fig. 6.**
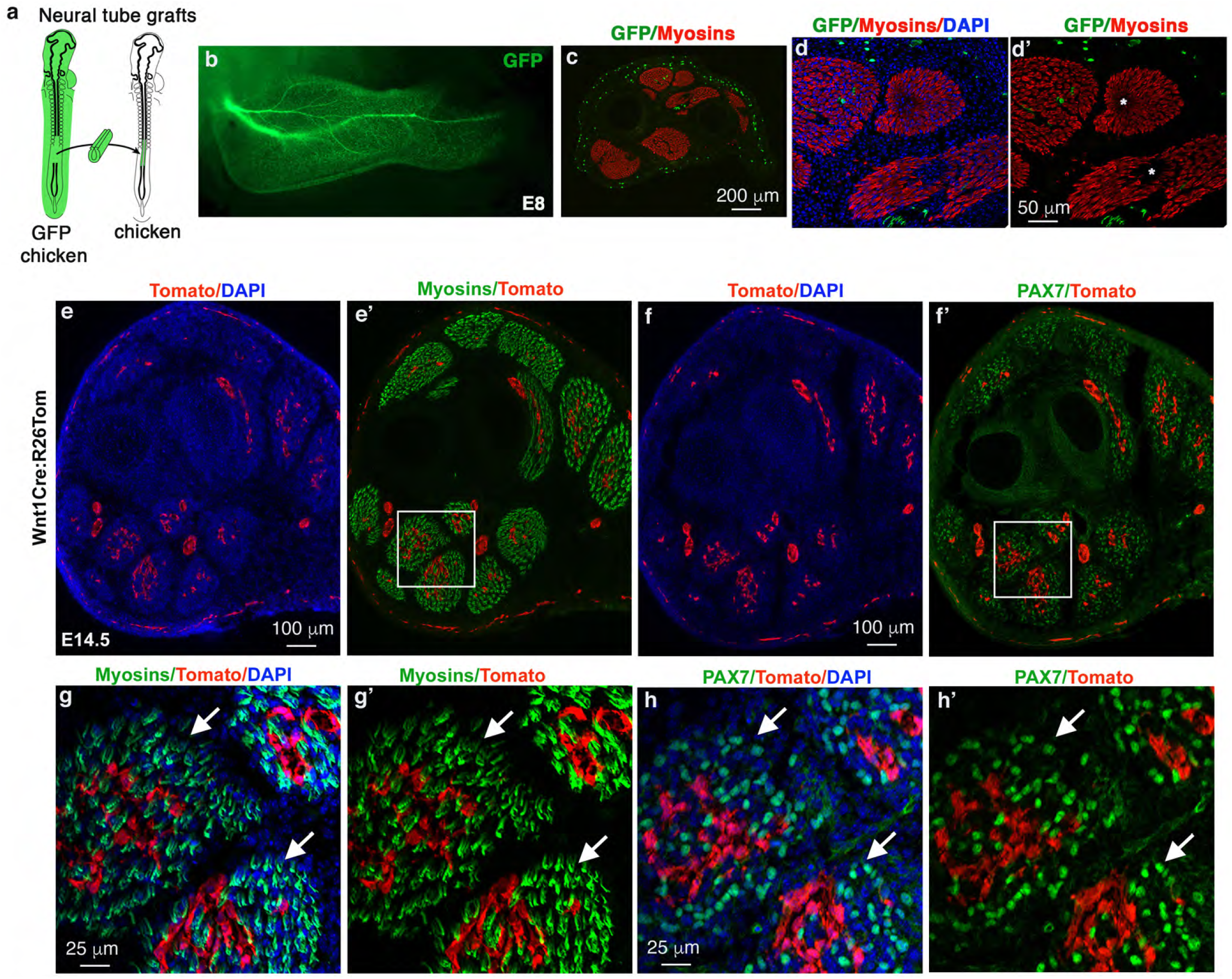
Neural crest cells do not contribute to myogenic lineage at muscle tips of limb skeletal muscles. (**a**) Diagram of isotopic neural tube grafts from GFP chicken to chicken embryos. (**b**) GFP fluorescence in E8 grafted-forelimbs (6 days after grafting) showing the neural tube cell derivatives. (**c**) Transverse forelimb sections from neural tube grafts immunostained with GFP (neural tube-derived cells) and MF20 (myosins) antibodies. (**d**,**d’**) Focus on dorsal limb muscles immunostained with GFP (green) and MF20 (myosins) antibodies, combined with DAPI. White stars indicate tendons. (**e,e’**) Transverse forelimb sections of E14.5 *Wnt1^Cre^:R26^Tom^* mice immunostained with Tomato (Wnt1-lineage-derived cells) and My32 (myosins) antibodies, combined with DAPI. (**f,f’**) Transverse forelimb sections of E14.5 *Wnt1^Cre^:R26^Tom^* mice immunostained with Tomato (Wnt1-lineage-derived cells) and PAX7 antibodies, combined with DAPI. (**g,g’), (h,h’)** are high magnifications of ventral limb regions squared in (e’) and (f’), respectively. Arrows point to muscle cells (Myosin+ or PAX7+ cells, green) at muscle tips displaying no Tomato labelling, (Wnt1-lineage-derived cells, red).

**Extended data Fig. 7.**
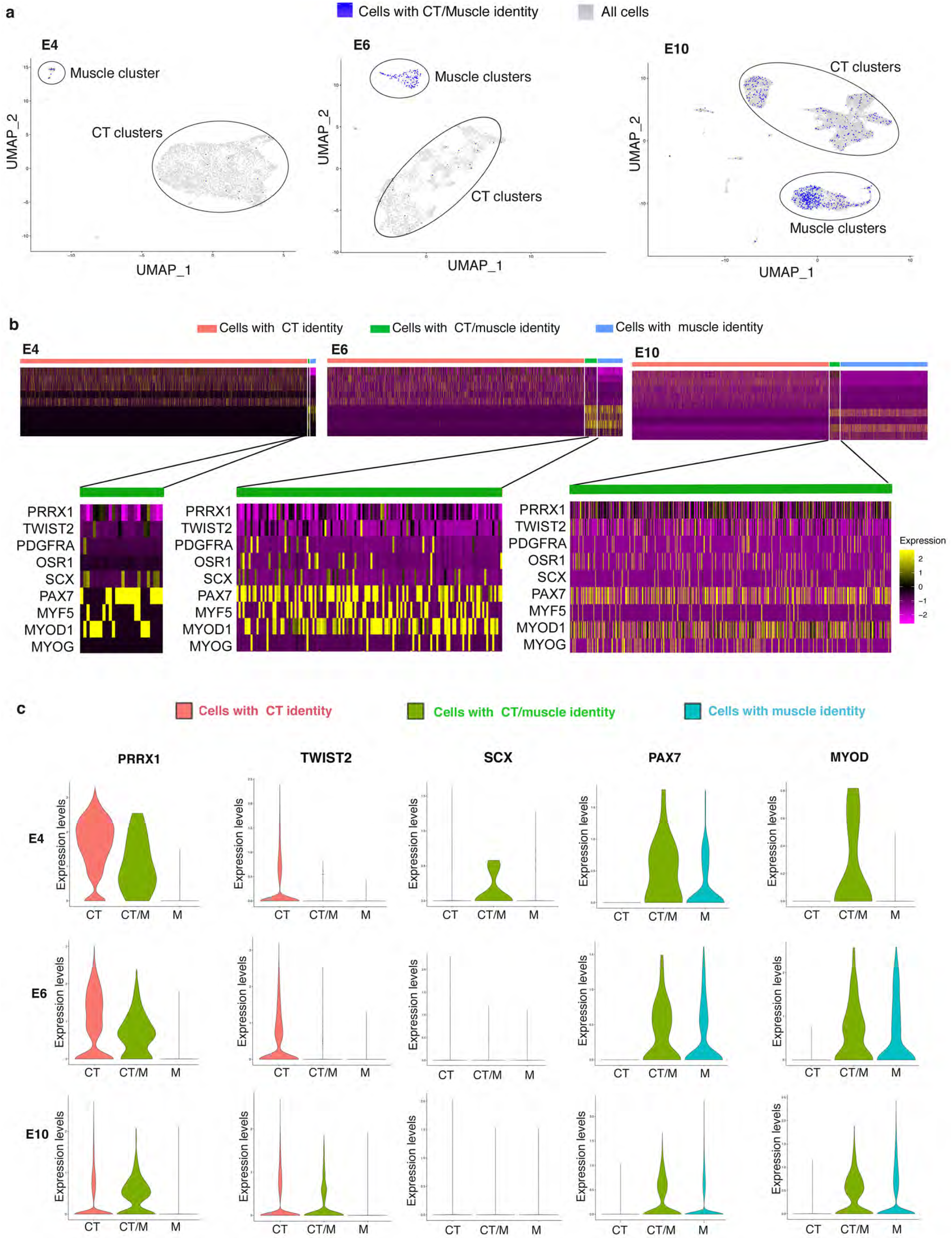
Characterisation of the cells with a CT/M identity. (**a**) Feature plots showing the distribution of cells with a CT/M identity (in blue) across whole-limb clustered populations (in grey) at E4, E6 and E10. At E4, 50% of the CT/M cells are found in the CT clusters and 50% in muscle clusters. At E6, 17% and 83% of the CT/M cells are found in CT and muscle clusters, respectively. At E10, 37% and 61% of the CT/M cells are found in CT and muscle clusters, respectively. (**b**) Heatmaps showing the relative expression of five recognized CT markers (PRRX1, TWIST2, PDFGRA, OSR1, SCX) and four muscle markers (PAX7, MYF5, MYOD, MYOG) in cells grouped by their identity CT (red), CT/M (green) or M (blue) at E4, E6 and E10. Upregulated genes in yellow, downregulated genes in purple. High magnification of the heatmaps showing the relative expression of the same nine CT and muscle markers but only in CT/M cells at E4, E6 and E10. (**c**) Violin plots showing Log-normalized expression levels of selected CT and muscle markers in cells grouped by their identity CT (red), CT/M (green) or M (blue) at E4, E6 and E10. The expression of PRRX1, and to a lesser extent TWIST2, is progressively found in an increasing number of CT/M cells at the expense of CT cells. Conversely, SCX expression is found in a decreasing number of CT/M cells, like PAX7 and MYOD.

**Extended Data Fig. 8.**
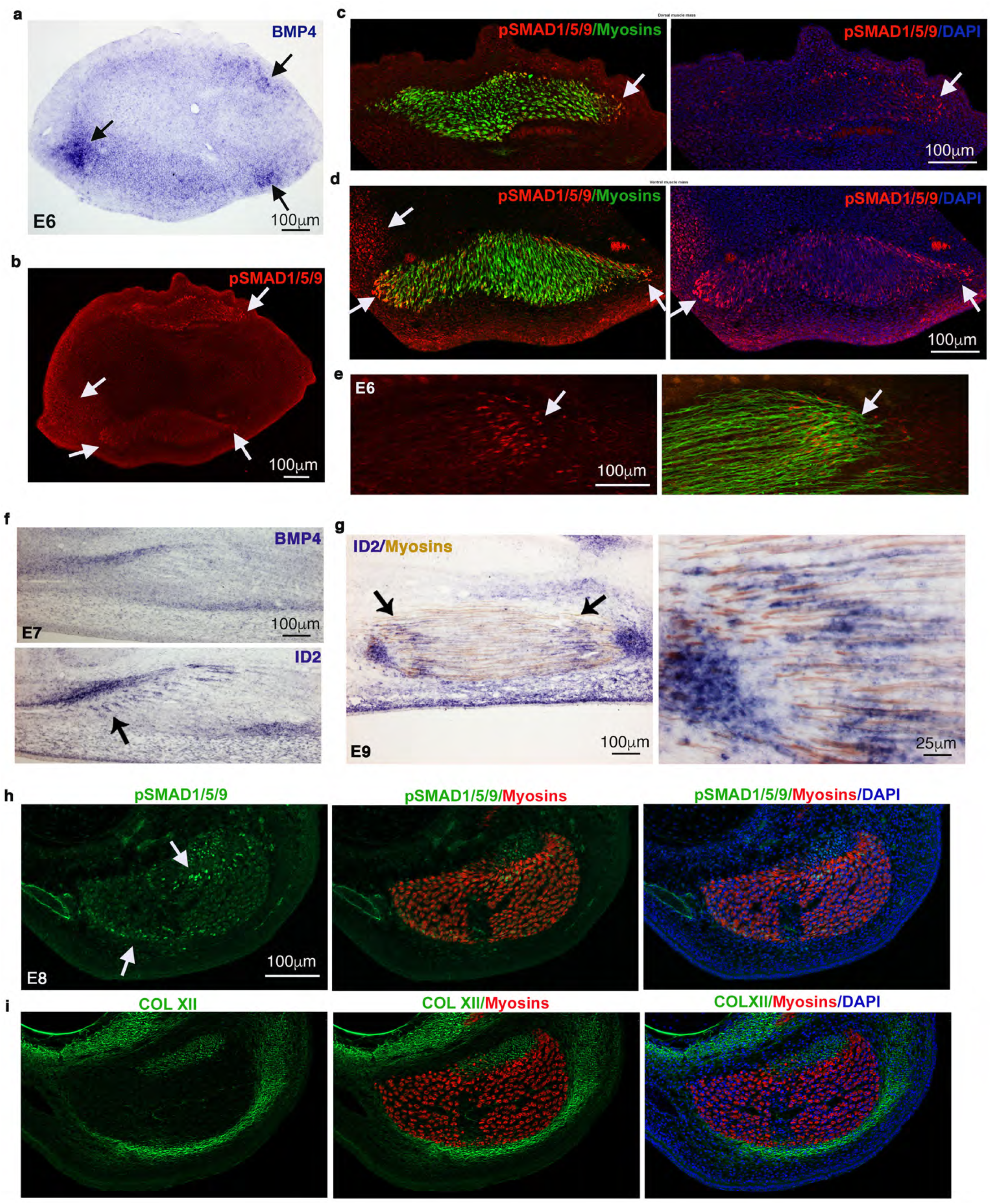
BMP signalling labels a junctional cell population at different stages of development. (**a,b**) Transverse adjacent limb sections of E6 chicken embryos were hybridized with BMP4 probe (a) and immunostained with pSMAD1/5/9 antibody (b). Arrows point to BMP4 and pSMAD1/5/9 expression spots. (**c,d**) High magnification of dorsal (c) and ventral (d) muscle masses immunostained with pSMAD1/5/9 and myosin antibodies and counterstained with DAPI. Arrows point to pSMAD1/5/9 expression spots. (**e**) Longitudinal view of a muscle labelled with pSMAD1/5/9 and MF20 (myosins) antibodies. Arrows point to pSMAD1/5/9 enriched expression (red) at muscle tips (green). **(f)** Adjacent and longitudinal limb sections of E7 chicken embryos were hybridized with BMP4 and ID2 probes (blue staining). The arrows point to *ID2* expression at muscle tips close to tendons labelled with *BMP4* and *ID2* transcript. (**g**) Longitudinal muscle sections of E9 chicken hybridized with ID2 probe (blue) and then immunostained with MF20 antibody (myosins, brown). The arrow points to *ID2* expression at muscle tips. (**h,i**) Adjacent and transverse limb sections of E8 chicken embryos were co-immunostained with pSMAD1/5/9 and MF20 (myosins) antibodies (h) and COLXII (tendons) and MF20 (myosins) antibodies (i). Arrows point to restricted location of pSMAD1/5/9 in FCU muscle

**Extended Data Fig. 9.**
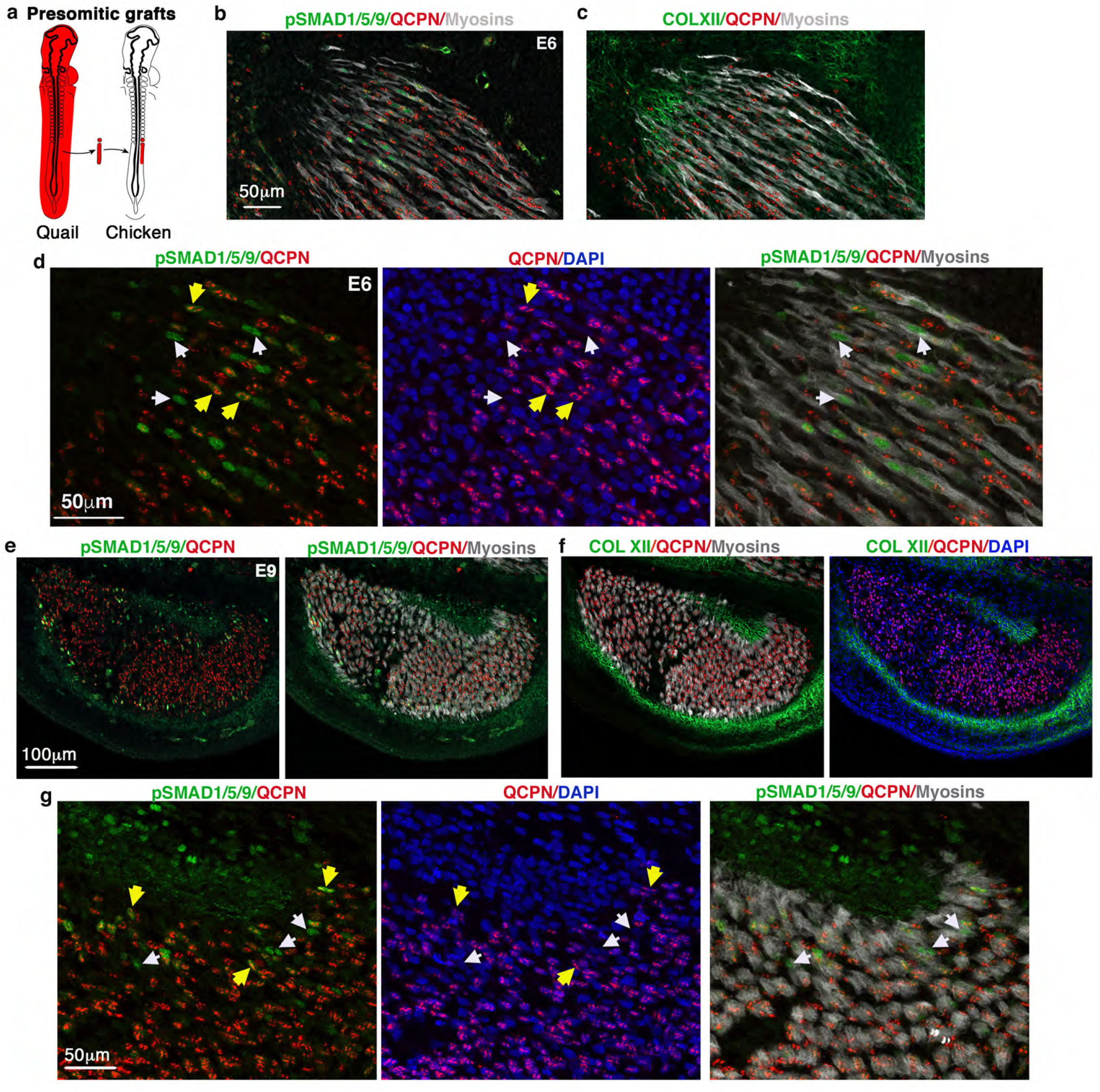
PSMAD1/5/9+ myonuclei label two myonucleus populations of distinct embryological origins at muscle tips. (**a**) Schematic of isotopic pre-somitic mesoderm grafts from quail to chicken embryos. (**b**) Longitudinal view of a limb muscle from E6 chimera labelled with pSMAD1/5/9, QCPN (quail nuclei) and MF20 (myosins) antibodies, with a focus to muscle tips. (**c**) Adjacent section labelled with COLXII (tendon), QCPN (quail nuclei) and MF20 (myosins) antibodies. (**d**) Zoom to muscle tips, labelled with pSMAD1/5/9, QCPN (quail nuclei) and MF20 (myosins) antibodies, combined with DAPI. White arrows point to pSMAD1/5/9+ myonuclei (green) that are not of quail origin (QCPN-), while yellow arrows point to pSMAD1/5/9+ myonuclei (green) that are of quail origin (QCPN+). (**e**) Transverse view of the FCU muscle from E9 chimera labelled with pSMAD1/5/9, QCPN (quail nuclei) and MF20 (myosins) antibodies. (**f**) Adjacent sections to (e) labelled with COLXII (to label tendon), QCPN (quail nuclei) and MF20 (myosins) antibodies, combined with DAPI. (**g**) Focus on muscle tips. White arrows point to pSMAD1/5/9+ myonuclei (green) that are not of quail origin (QCPN-), while yellow arrows point to pSMAD1/5/9+ myonuclei (green) that are of quail origin (QCPN+).

**Extended Data Fig. 10.**
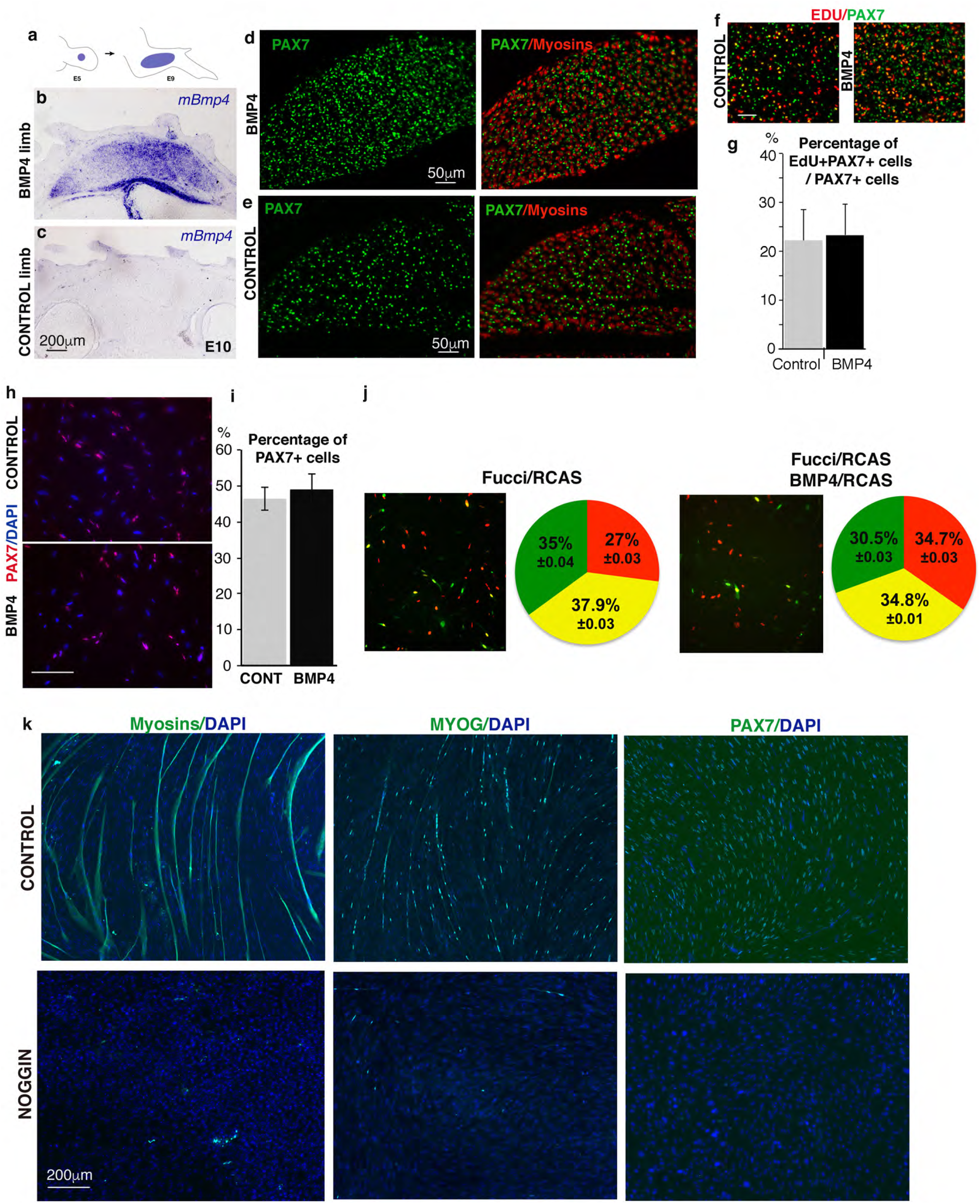
BMP signalling and fibroblast/myoblast conversion. (**a**) Schematics of the grafting procedure of BMP4/RCAS-producing cells. (**b,c**) In situ hybridisation with BMP4 probe to transverse limb sections indicate the extent of ectopic BMP4, in E10 grafted-embryos, 5 days after grafting (BMP4 right (b) and control left (c) limbs). (**d,e**) Immunohistology with PAX7 (green) and MF20 (myosins, red) antibodies to BMP4 right (d) and control left (e) limb sections. (**f,g**) Following BMP4 exposure, the proportion of EDU+/PAX7+ cells versus PAX7+ cells was not significantly changed compared to control muscles. **(h-j)** BMP4/RCAS exposure to chicken primary myoblast cultures in proliferation conditions did not increase the number of PAX7+ cells compared to control (empty/RCAS) myoblast cultures (h,i) and did not affect cell proliferation assed with the Fucci system (j). (**k**) NOGGIN/RCAS and empty/RCAS (control) exposure to chicken primary myoblast cultures in differentiation conditions. Myoblasts were immunostained with MF20 (myosins), MYOG and PAX7 antibodies (green) combined with DAPI. NOGGIN/RCAS exposure in myoblast cultures led to a drastic disappearance of Myosin+ myotubes, MYOG+ cells and PAX7+ cells compared to control cultures, while DAPI nuclei were still observed.

